# Cyclin B2 is required for progression through meiosis in mouse oocytes

**DOI:** 10.1101/441352

**Authors:** Enrico Maria Daldello, Xuan G. Luong, Cai-Rong Yang, Jonathan Kuhn, Marco Conti

## Abstract

Cyclins associate with CDK1 to generate the M-phase-promoting factor (MPF) essential for progression through mitosis and meiosis. Previous studies concluded that CCNB2 is dispensable for cell cycle progression. Given our findings of high translation rates of *CcnB2* mRNA in prophase-arrested oocytes, we have reevaluated its role during meiosis. *CcnB2*^−/−^ oocytes undergo delayed germinal vesicle breakdown followed by a defective M-phase due to reduced pre-MPF activity. This disrupted maturation is associated with compromised *CcnB1* and *Mos* mRNA translation and delayed spindle assembly. Given these defects, a significant population of oocytes fail to complete meiosis I because SAC remains activated and APC function is inhibited. *In vivo*, CCNB2 depletion leads to decreased oocyte developmental competence, compromised fecundity, and premature ovarian failure. These findings demonstrate that CCNB2 is required to assemble sufficient pre-MPF for timely meiosis reentry and progression. Although endogenous cyclins cannot compensate, overexpression of CCNB1 rescues the meiotic phenotypes, demonstrating similar molecular properties but divergent modes of regulation of these cyclins.

## INTRODUCTION

Successful completion of the two meiotic cell divisions is essential for gamete development and fertility. Fully-grown oocyte reentry into meiosis requires assembly of the M-phase-promoting factor (MPF) and activation of its kinase activity (Adhikari and Liu, 2014). This complex subsequently phosphorylates a large number of protein substrates triggering dissolution of the nuclear membrane, chromosome condensation, and spindle assembly (Morgan, 2007). Once proper chromosome-to-microtubule attachment is achieved, rapid inactivation of MPF is necessary for the transition to anaphase. This inactivation depends on the switch-like activation of anaphase-promoting complex/cyclosome (APC/C), followed by ubiquitination and degradation of cyclins and, therefore, inactivation of MPF (Thornton and Toczyski, 2006; Yang and Ferrell, 2013). Concomitant degradation of securin leads to activation of separase, which cleaves cohesins, allowing separation of the bivalents in anaphase (Lane et al., 2012). Given the asymmetrical position of the spindle in oocytes, telophase results in the extrusion of a small polar body.

The MPF is composed of two classes of molecules that orchestrate progression through both M-phases of mitosis and meiosis: a family of cyclin-dependent serine/threonine kinases (CDKs) and their binding partners, cyclins (Morgan, 2007). While there are three M-phase *CcnB* mRNAs present in mammals (*B1*, *B2*, and *B3*), most of the molecular properties of the CDK1/CCNB complex are based on observations of the CDK1/CCNB1 heterodimer. However, other cyclins, like CCNB2, also interact with CDK1, activating their phosphotransferase activity (Jackman et al., 1995). CCNB1 and B2 are thought to be localized in different subcellular compartments (Jackman et al., 1995). While CCNB1 is either soluble or interacts with microtubules, CCNB2 is associated with the cellular membrane. During mitosis, CCNB1 translocates into the nucleus while CCNB2 remains sequestered in the cytoplasm (Jackman et al., 1995).

Although all three cyclin mRNAs are detected in mouse oocytes, the CCNB1/CDK1 complex is generally regarded as the major driver of meiosis progression. Little is known about the role of CCNB3 during oocyte maturation with the exception of one report that suggests its requirement in meiosis I (Zhang et al., 2015). Conversely, CCNB2 is thought to be dispensable for mitosis progression. Early attempts to define CCNB2 function either by genetic inactivation of the gene or by knockdown with antisense RNAs have not produced overt phenotypes; this has led to the conclusion that CCNB2 is dispensable for either mitosis and meiosis, possibly due to compensation by CCNB1 (Brandeis et al., 1998; Ledan et al., 2001). However, recent evidence reproposes an independent function for CCNB2 during the mouse meiotic divisions. During experiments investigating the function of the spindle component NDC80/HEC1 during meiotic prophase, it has been proposed that CCNB2 stability is dependent on association with HEC1 (Gui and Homer, 2013). Additionally, knockdown of CCNB2 with morpholino oligonucleotides (MO) markedly decreased meiotic reentry in mouse oocytes (Gui and Homer, 2013). Similar findings have been reported in a very recent study investigating CCNB1 function in oocytes (Li et al., 2018). New data from our laboratory has demonstrated that the rate of translation of the two major CCNBs, B1 and B2, is regulated in a distinct fashion during meiotic prophase (Han et al., 2017). *CcnB1* mRNA is expressed with three distinct 3’UTRs of different lengths; while translation of the two longer mRNA variants is repressed in meiotic prophase, a third, short variant is constitutively translated (Yang et al., 2017). CCNB1 protein is detectable in meiotic prophase albeit at levels that are low compared to M-phase. During prometaphase, the translation of the two longer variants is activated and drives the large accumulation of the CCNB1 protein (Yang et al., 2017). Although the two *CcnB2* mRNA variants are also detected, the rate of translation of *CcnB2* mRNA is high in prophase I and the protein is readily detectable—a feature reminiscent of that reported in frog oocytes (Piqué et al., 2008).

The above findings open the possibility that CCNB2 is considerably more abundant than CCNB1 in prophase I (GV)-arrested oocytes. Prompted by these observations, we have further investigate the function of CCNB2 during mouse oocyte meiotic progression. Using previously generated *Ccnb*2^−/−^ mice (Brandeis et al., 1998), we show that oocytes deficient in CCNB2 are developmentally compromised, as documented by the subfertility of these mice. This sub-fertility phenotype is due to inefficient oocyte reentry and progression through the meiotic cell cycle with blocks at different stages of meiosis. Thus, we conclude that CCNB2 plays a significant role during mouse oocyte meiosis, which cannot be compensated for by endogenous CCNB1.

## RESULTS

### Contrasting CcnB1 and CcnB2 mRNA translation rates define the pattern of expression of the two cyclins at the prophase I to metaphase I transition

We have previously reported that the patterns of ribosome loading onto the *CcnB1* and *CcnB2* mRNAs in fully-grown mouse oocytes are considerably different (Han et al., 2017). Here, we confirmed and extended this initial observation with a detailed time course experiment monitoring ribosome loading onto the two mRNAs in GV oocytes and during progression through metaphase I (MI) (Fig. 1, A and B). The overall mRNA levels for the two cyclins are comparable (Input, Fig. 1, A and B). However, while little ribosome loading onto *CcnB1* mRNA is detected in GV oocytes, ribosome loading onto *CcnB2* mRNA is considerably higher. These indirect measurements of translation are corroborated by mining data sets assessing poly(A)-tail length of mRNAs in GV oocytes (Morgan et al., 2017). The *CcnB2* mRNA has a significantly longer poly(A)-tail as compared to *CcnB1* (Fig. 1 C); increased poly(A)-tail length has been associated with an increased rate of translation (Reyes and Ross, 2016).

**Fig. 1.**
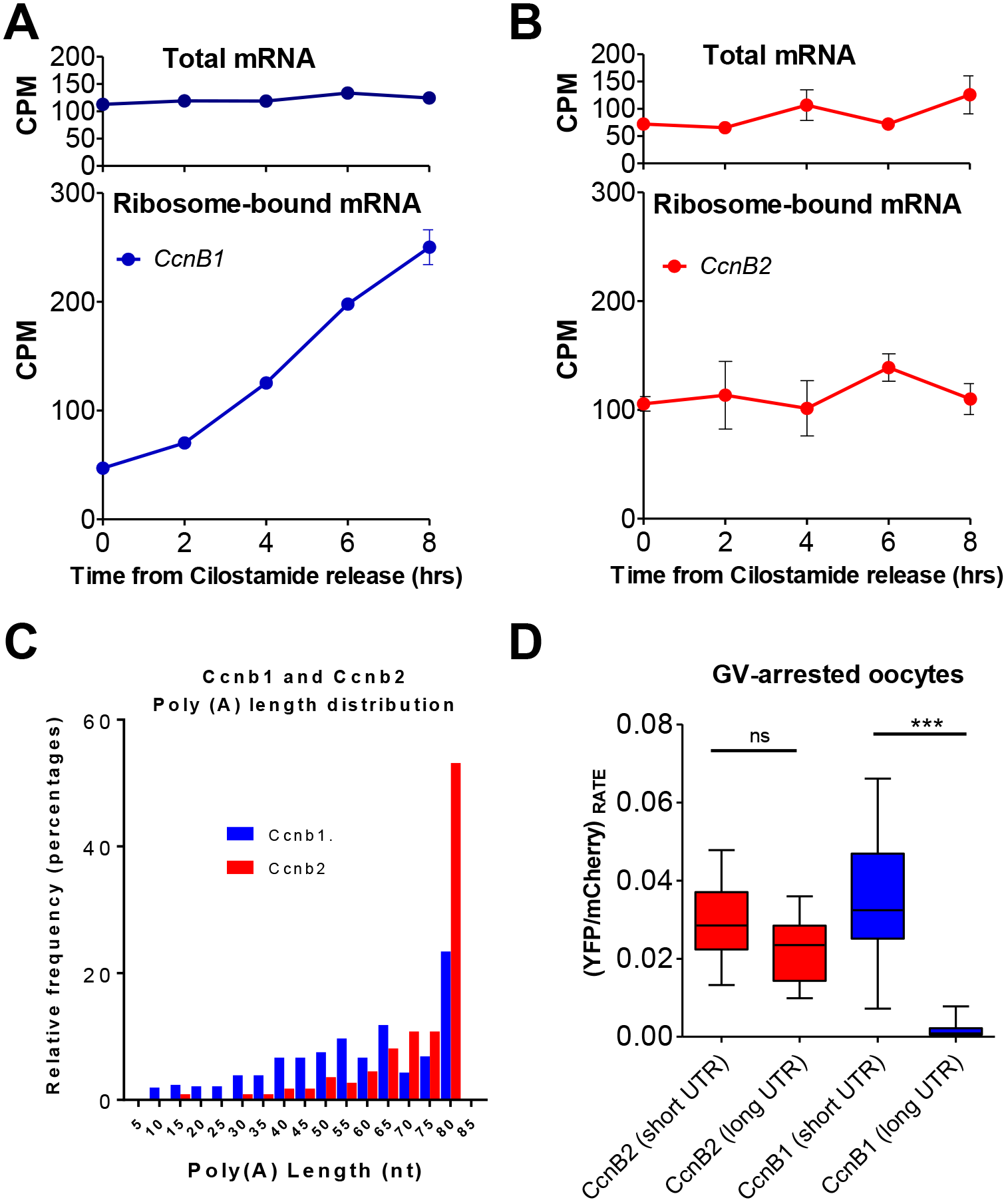
Translation of *CcnB1* and *CcnB2* mRNAs is differentially regulated during meiotic maturation in mouse oocytes. **A-B)** RNA-Seq was performed using mRNA extracts of cell lysate (total mRNA) or mRNA extracts after immunoprecipitation of HA-tagged ribosomes (ribosome-bound mRNA) from oocytes arrested in prophase with Cilostamide (time 0) or collected 2, 4, 6, and 8 hrs after meiosis resumption. Counts per million (CPM) mapped reads are reported for *CcnB1* **(A)** and *CcnB2* **(B)**; average CPMs of two independent biological replicates with range are reported. **C)** Poly(A) tail lengths of the *CcnB1* and *CcnB2* mRNAs in GV oocytes. The data were mined from PMID: 28792939 and are reported as binned values up to 80 (A) nucleotides. **D)** Rates of translation of *CcnB1* and *CcnB2* mRNA variants in prophase I. Oocytes were injected with 1:1 mix of YFP-oligo-adenylated 3’UTR (*CcnB2*-short, *CcnB2*-long, *CcnB1*-short, or *CcnB1*-long) and polyadenylated *mCherry*. Rate of translation in GV-arrested oocytes were calculated with a 3 hr window at a sampling rate of 15 mins. T-tests were performed for statistical significance; “ns”: not significant, “***”: p< 0.0001.

During meiotic progression, little or no changes in ribosome loading onto the *CcnB2* mRNA were detected up to MI, whereas major changes in *CcnB1* association with ribosomes take place during MI (Fig. 1 A). This differential pattern of translation is in good accordance with data from previous experiments using luciferase reporters tagged with the 3’UTRs of the two mRNAs (Han et al., 2017). We also have shown that alternate polyadenylation signal usage (APA) plays a major role in defining the 3’UTR length and translation rate of *CcnB1* mRNA (Yang et al., 2017). Since the *CcnB2* mRNA 3’UTR also contains an internal polyadenylation signal, we compared the translation rate of the two 3’UTR variants. *CcnB1* 3’UTR short and long constructs were used as a control. The rates of translation of the two *CcnB2* 3’UTR reporters are comparable in GV oocytes (Fig. 1 D) and do not change significantly during oocyte maturation. However, the rate of translation driven by the long *CcnB1* 3’UTR is considerably lower than that of either *CcnB2* 3’UTRs (Fig. 1 D). Only the rate of translation of the short *CcnB1* 3’UTR approximates those of *CcnB2* 3’UTR. These findings consolidate the concept that rates of translation of the two cyclin mRNAs are significantly different. Since previous experiments indicate comparable degradation rates of the two proteins in GV (Han et al., 2017), we hypothesize that CCNB2 accumulates in GV-arrested oocytes at higher levels than CCNB1. During oocyte maturation, CCNB1 accumulation increases while CCNB2 remains relatively unchanged, opening the possibility of a shift in the stoichiometry of CCNB/CDK1 complex.

### CcnB2^−/−^ female mice display defects in fecundity

Together with a previous report (Gui and Homer, 2013), the above findings are at odds with the widely held notion that CCNB2 is dispensable for oocyte maturation. Therefore, we have re-evaluated the fertility phenotype of the previously generated *CcnB*2^−/−^ mice (Brandeis et al., 1998).

While both CcnB2^+/−^ and *CcnB*2^−/−^ mice are fertile and produce pups, *CcnB*2^−/−^ mice show compromised fecundity (Fig. 2 A). This phenotype is not due to embryonic lethality of the *CcnB*2^−/−^ pups since there is no statistically significant deviation from the expected Mendelian ratio when heterozygous males and females were mated—suggesting that CCNB2 is dispensable for embryo development (Fig. S1 A). Furthermore, *CcnB*2^−/−^ females generate fewer pups even when crossed with wild type (WT) males, indicating that the sub-fertility is associated with the female. *CcnB*2−/− females gave birth to fewer pups (reduced litter size) and fecundity declined rapidly with age, suggesting premature ovarian failure (Fig. 2 A and Fig. S1, B and C). This fecundity phenotype may be caused by delayed puberty and/or ovarian or oocyte dysfunction. Delayed puberty is ruled out as the pregnancy rates of mated *CcnB*2^−/−^ females did not improve over a period of nine months. Furthermore, the age at first pregnancy of *CcnB*2^+/+^ and *CcnB*2^−/−^ females are comparable (Fig. S1 D). Consistent with the original report (Brandeis et al., 1998), while *CcnB*2^+/−^ pups are indistinguishable from the *CcnB*2^+/+^ littermates, the *CcnB*2^−/−^ pups were significantly smaller, weighing an average 1.4 ± 0.35 g less than the *CcnB2^+/+^* siblings (Fig. 2 B). The adult ovarian morphology of the *CcnB*2^−/−^ mice is unremarkable, with all the follicle developmental stages and corpora lutea present (Fig. 2 C). To further assess ovarian function and monitor follicle maturation/ovulation potential, pre-pubertal females were injected with PMSG followed by hCG, a regimen which induces ovulation. Slightly fewer oocytes were retrieved from *CcnB*2^−/−^ females, while oocyte diameters are identical between *CcnB*2^+/+^ and *CcnB*2^−/−^ females (Supplemental Fig. S1, E and F).

**Fig. 2.**
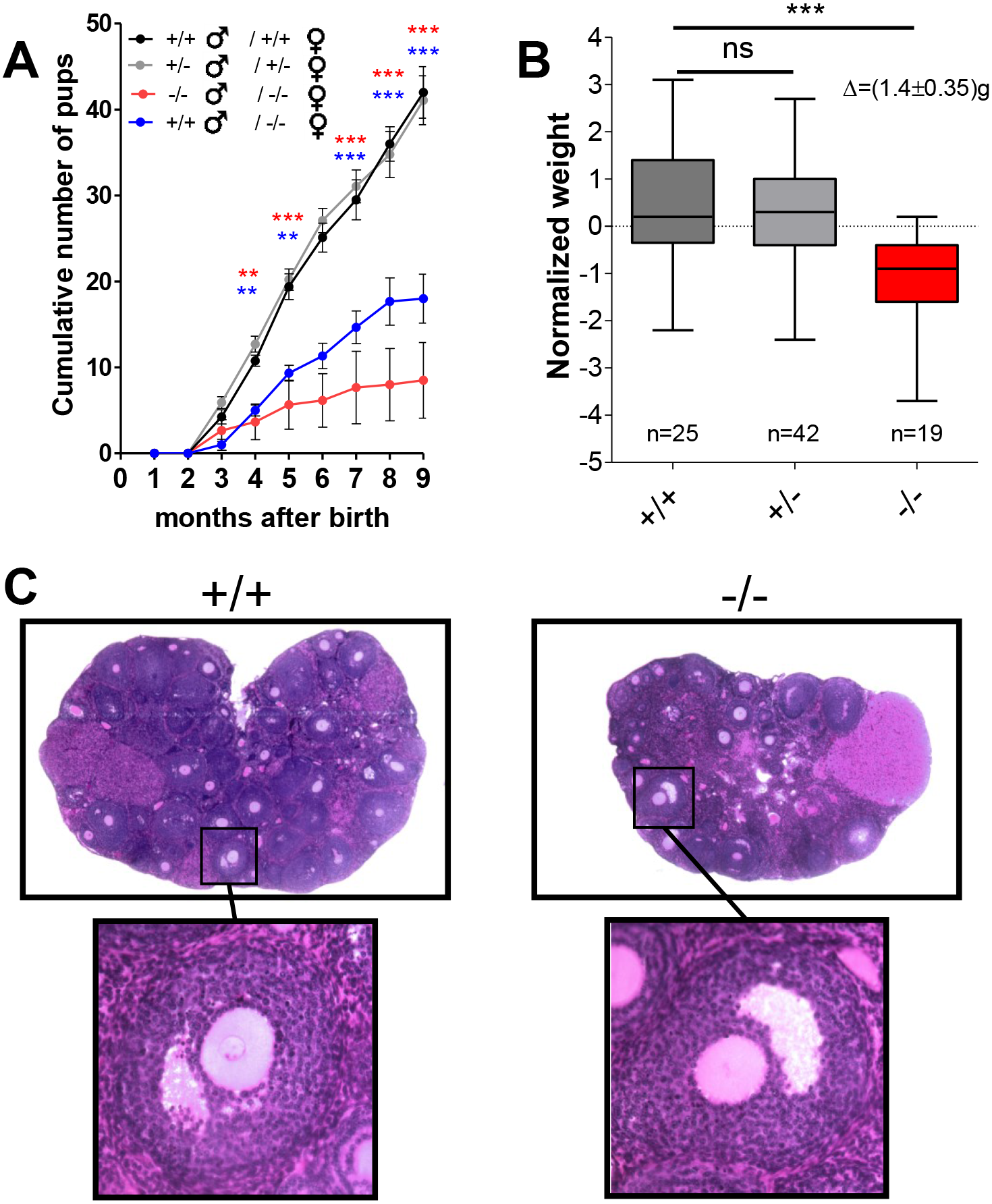
Compromised fecundity of the *CcnB*2_−/−_ mice. **A)** Cumulative number of pups per female derived from different mating schemes. Mating schemes and number of pairs were as follows: +/+♂ × +/+♀, n= 20; +/−♂ × +/−♀, n= 35; − /−♂ × −/−♀, n= 6; +/+♂ × −/−♀, n= 6. T-tests were performed between +/−♂ × +/−♀ and −/−♂ × −/−♀ (red asterisks) or +/− ♂ × +/−♀ and +/+ ♂ × −/−♀ (blue asterisks); “**”: p< 0.01, “***”: p< 0.001. Breeding was initiated when the mice reached four weeks of age. **B)** Pup body weights from +/−♂ × +/−♀ matings were recorded 21 days after birth. The weight of each mouse was normalized for the average weight of the litter and plotted according to their genotype. T-tests were performed for statistical significance; “ns”: not significant, “***”: p< 0.0002. **C)** Representative 8.0 μm histological H&E staining sections of ovaries from *CcnB*2^+/+^ and *CcnB2r*^−/−^ mice.

### Timing of oocyte maturation is aberrant in CcnB2^−/−^ oocytes

Western blot analysis of *CcnB*2^+/−^ and *CcnB*2^−/−^ oocyte extracts reveal a gene dose-dependent decrease of CCNB2 protein (Fig. 3 A). Loss of CCNB2 does not affect CDK1 protein levels, indicating no effect on either synthesis or stability of the kinase moiety. Recently, it was reported that oocyte-specific knockout of CCNB1 results in the overexpression of CCNB2 (Li et al., 2018); however, in *CcnB*2^−/−^ oocytes, the level of CCNB1 is not obviously altered (Fig. S2 A). To understand the extent by which CDK1 activity depends on CCNB2, we measured the CDK1 kinase activity using two independent strategies. In the first paradigm, extracts from *CcnB*2^+/+^ and *CcnB*2^−/−^ oocytes were incubated with a CDK1 substrate (GST-PP1) and phosphorylation was measured by phosphosite-specific antibodies (pT320-PP1) (Daldello et al., 2015; Lewis et al., 2013). There is a highly significant decrease in CDK1 activity in extracts from *CcnB*2^−/−^ oocytes (Fig. 3, B and C). MPF activity was also measured in whole oocytes using a previously described CDK1-FRET reporter assay (Gavet and Pines, 2010a; Gavet and Pines, 2010b). In GV-arrested *CcnB*2^−/−^ oocytes, FRET signal is decreased as compared to maturing *CcnB*2^+/+^ oocytes (Fig. S2 B).

Given that the above data are consistent with decreased pre-MPF activity, we investigated whether spontaneous maturation is affected in *CcnB*2^−/−^ oocytes. While *CcnB*2^+/+^ and *CcnB*2^+/−^ oocytes resume meiosis in a highly synchronous manner (GVBD time= 1.5 ±1.1 hrs and 1.8 ±1.1 hrs, respectively), meiotic reentry of *CcnB*2^−/−^ oocytes is significantly delayed (GVBD time= 4.3 ±3.7 hrs). A more detailed analysis of the maturation time course shows the presence of two subpopulations of *CcnB*2^−/−^ oocytes. The first population resumes meiosis within the first four hours post-Cilostamide release, though the GVBD time is still delayed compared to *CcnB*2^+/+^ oocytes. The second population resumes meiosis in a stochastic manner, with oocytes undergoing GVBD even after 16 hours post-Cilostamide release (Fig. 3 D and Fig. S2 C). Furthermore, the time of GVBD in *CcnB*2^−/−^ oocytes is inversely correlated with CDK1 activity of the same oocytes at GV (Fig. S2 D), but there is no correlation between GVBD time and oocyte diameter (Fig. S2 E). If a decreased MPF activity were solely responsible for delayed meiosis resumption in these oocytes, the overexpression of cyclins should rescue the phenotype. Indeed, overexpression of *CcnB1-mCherry* in *CcnB*2^−/−^ oocytes restores the GVBD time to that of *CcnB*2^+/+^ oocytes (Fig. 3E), and oocytes expressing higher levels of CCNB1 undergo GVBD earlier (Fig. S2 F). Of note, CCNB1-mCherry, but also CCNB2-mCherry, translocate into the nucleus with the same kinetics (Fig. S2 G). These findings strongly support the hypothesis that CCNB2 protein accumulation in prophase I is required to generate sufficient CDK activity for timely reentry into meiosis, and that oocytes with lower pre-MPF activity resume meiosis in a delayed fashion.

**Fig. 3.**
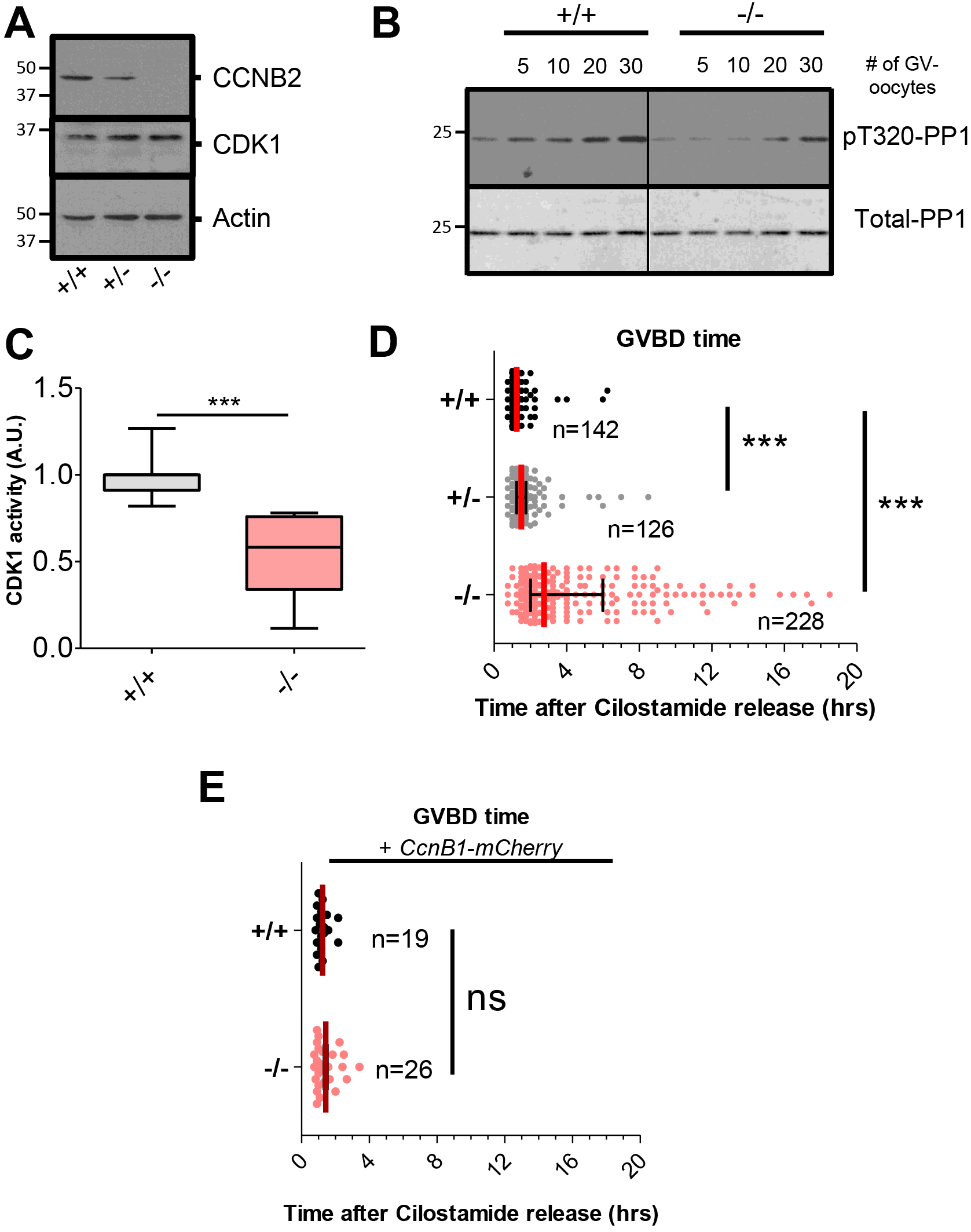
Aberrant timing of meiotic resumption in oocyte depleted of CCNB2 is due to defective pre-MPF. **A)** Western blot analysis of extracts from 150 oocytes from *CcnB*2^+/+^, ^+/−^, and ^−/−^ mice. **B)** Kinase assays were performed using increasing numbers of oocytes from *CcnB*2^+l+^ (+/+) or *CcnB*2^−/−^ (−/−) mice and a GST-pp1 fragment as a substrate. Levels of T320 PP1 phosphorylation were detected using a specific antibody (pT320-pp1). The level of total substrate loaded was evaluated by Ponceau S staining (Total-pp1). **C)** Quantification of six independent kinase assays. pT320-pp1/Total-pp1 ratios from *CcnB*2^−/−^ oocytes were expressed as fold changes over their matched *CcnB*2^+/+^ controls. A t-test was performed to determine statistical significance; “***”: p= 0.0007. **D)** Time of GVBD was determined through brightfield images acquired every 15 mins for 24 hrs. Results from four independent experiments are included. Median times of GVBD with interquartile range are reported (mean GVBD time: *CcnB*2^+/+^= (1.35 ± 0.06 hrs), *CcnB*2^+/−^= (1.76 ± 0.10 hrs), *CcnB*2^−/−^= (4.41 ± 0.24 hrs). A non-parametric Mann-Whitney test was performed to evaluate statistical significance; “***”: p< 0.0001. **E)** Oocytes were injected with mRNA encoding *CcnB1-mCherry* and, after 3h incubation, were released in Cilostamide-free medium. GVBD time and statistical significance were determined as in **D)**; “ns”: not significant.

### PKA inactivation-dependent events are intact whereas CDK1-dependent events are disrupted in CcnB2^−/−^ oocytes

In order to further define the molecular defects associated with CCNB2 depletion in the oocyte during the G_2_/M transition, we examined the timing of CDC25B translocation. We have previously shown that in mouse oocytes, CDC25B import into the nucleus is one of the first detectable events following the decrease in cAMP, the signal that maintains oocyte meiotic arrest (Oh et al., 2010). Therefore, we injected oocytes with *CDC25B*-(phosphatase dead)-*YFP* to follow the kinetics of CDC25B translocation in intact oocytes (Fig. 4 A). All *CcnB*2^+/+^ oocytes mature in a synchronous manner (Fig. S3 A) and the YFP-tagged CDC25B signal is detected in the nucleus at as early as 15 mins post-Cilostamide release (Fig. S3 B). However, there are two populations of *CcnB*2^−/−^ oocytes (early GVBD and late GVBD) (Fig. S3 E). CDC25B translocation in both *CcnB*2^+/+^ and *CcnB*2^−/−^ oocytes continues until the oocytes undergo GVBD, at which point CDC25B diffuses throughout the cytoplasm (Fig. 4 A). No significant difference in CDC25B translocation rate are found between *CcnB*2^+/+^ oocytes and the two *CcnB*2^−/−^ populations (Fig. 4 C). However, since GVBD time is delayed in *CcnB*2^−/−^ oocytes, CDC25B accumulation in the nucleus continues for longer periods of time, resulting in higher CDC25B reporter signal in the nucleus (Fig. S3 C). This difference in the CDC25B nuclear/cytoplasmic ratio is not due to differences in the amount of reporter expressed (Fig. S3 D). These measurements document that CDC25B translocation occurs normally in *CcnB2*^−/−^ oocytes and that the rate of import is not affected by the decrease in MPF activity. Moreover, they indicate that PKA downregulation occurs normally in the *CcnB*2^−/−^ oocytes. Remarkably, these findings also demonstrate that CDC25B translocation alone was not sufficient to trigger GVBD in the *CcnB*2^−/−^ oocytes.

**Fig. 4.**
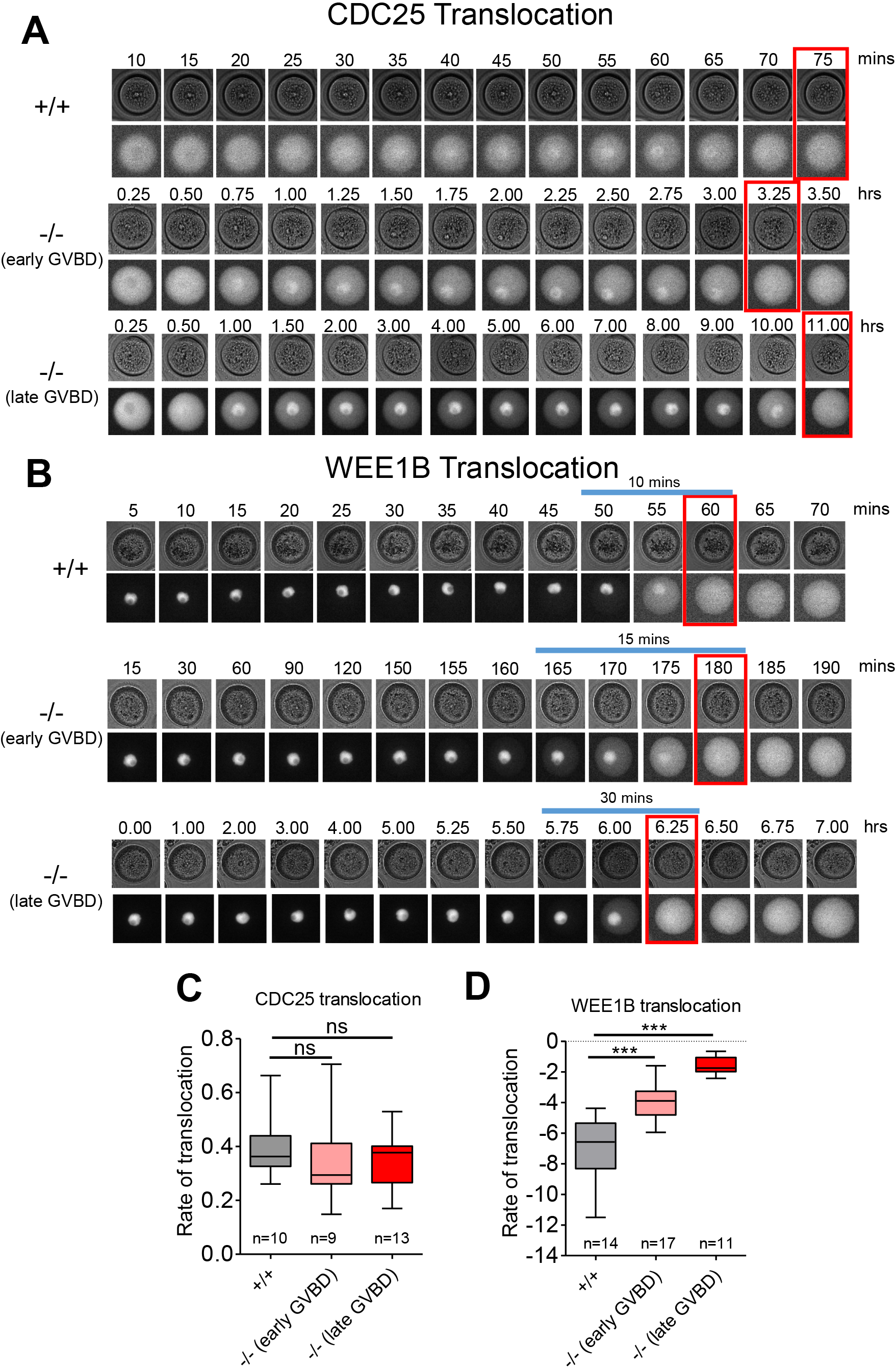
Although CDC25 translocation to the nucleus is unaffected, WEE1B export from the nucleus is delayed in *CcnB*2^−/−^ oocytes. Oocytes were injected with inactive *Cdc25B-YFP* **(A)** or *Wee1B-YFP* **(B)** and, after overnight incubation, were released in Cilostamide-free medium. Brightfield and YFP images were acquired every 5 mins for 20 hrs. Oocytes from *CcnB*2^−/−^ mice were divided into two groups according to their GVBD time; 0-4 hrs: “early GVBD” and ≥4 hrs: “late GVBD.” **A-B)** Representative pictures of an oocyte from *CcnB*2^+/+^, *CcnB*2^−/−^ (early GVBD), and *CcnB*2^−/−^ (late GVBD) are reported. The red box marks the time of GVBD. **C)** Rates of CDC25-YFP or **(D)** WEE1B-YFP translocation were calculated from each single oocyte as the slope of the linear regression of the Nuclear/Cytoplasmic ratios. Rates were expressed as medians with interquartile range. T-tests were performed to assess statistical significance; “ns”: not significant. “***”: p< 0.0001

CDC25B translocation is followed by CCNB1 import into the nucleus and WEE1B export out of the nucleus preceding GVBD (Oh et al., 2010). Effects on CCNB1 import could not be measured in *CcnB*2^−/−^ oocytes because of its rescuing effect (see below). However, YFP-tagged WEE1B export from the nucleus occurs over a wide range of time, consistent with the variable timing of GVBD (Fig. 4 B). *CcnB*2^−/−^ oocytes show significantly decreased WEE1B translocation rates as compared to *CcnB*2^+/+^ oocytes (Fig. 4 D); WEE1B translocation rate is inversely correlated to GVBD time (Fig. S3 F). We have previously demonstrated that WEE1B export is dependent on CDK1 activity (Oh et al., 2010); therefore, the decreased WEE1B export rate observed in *CcnB*2^−/−^ oocytes indicates slower CDK1 activation. Furthermore, we tracked CDK1 activation in live oocytes using a FRET approach to confirm that the speed of CDK1 activation is reduced in *CcnB*2^−/−^ oocytes. The speed of CDK1 activation, measured by the Hill slope of FRET activation before GVBD, is significantly decreased in *CcnB*2^+/−^ and *CcnB*2^−/−^ oocytes (Fig. S3, G and H). These results indicate that, in the absence of CCNB2, CDK1 activation is no longer switch-like but becomes gradual, resulting in inefficient WEE1B export and delayed GVBD.

### Translation of key cell cycle components is defective in CcnB2^−/−^ oocytes

Consistent with findings in *Xenopus* oocytes, we have previously shown that, at least in part, the translational program in mouse oocytes is dependent on CDK1 activity (Ballantyne et al., 1997; Han et al., 2017). Since CDK1 activation is likely defective in *CcnB*2^−/−^ oocytes, we tested whether CDK1-dependent translation would also be affected in these oocytes. Oocytes were co-injected with mRNA coding for *mCherry* (loading control) and an YFP reporter fused to either *CcnB1*-long 3’UTR or *Mos* 3’UTR. The accumulation of YFP and mCherry for individual oocytes was recorded throughout meiotic maturation and signals were expressed as ratios of YFP/mCherry (Fig. S4). The rates of translation were calculated for before (0-2 hrs) and after (4-8 hrs) GVBD for *YFP-CcnB1*-long 3’UTR (Fig. 5 A) and *YFP-Mos* 3’UTR (Fig. 5 B). As previously shown, the translation of both *YFP-CcnB1*-long 3’UTR and *YFP-Mos* 3’UTR increases during meiosis in *CcnB*2^+/+^ oocytes. In the absence of CCNB2, the translational activation varies widely with a population of oocytes showing a protein synthesis pattern similar to that of *CcnB*2^+/+^ oocytes and a population in which translation activation is absent or reduced for both reporters (Fig. 5 A).

**Fig. 5.**
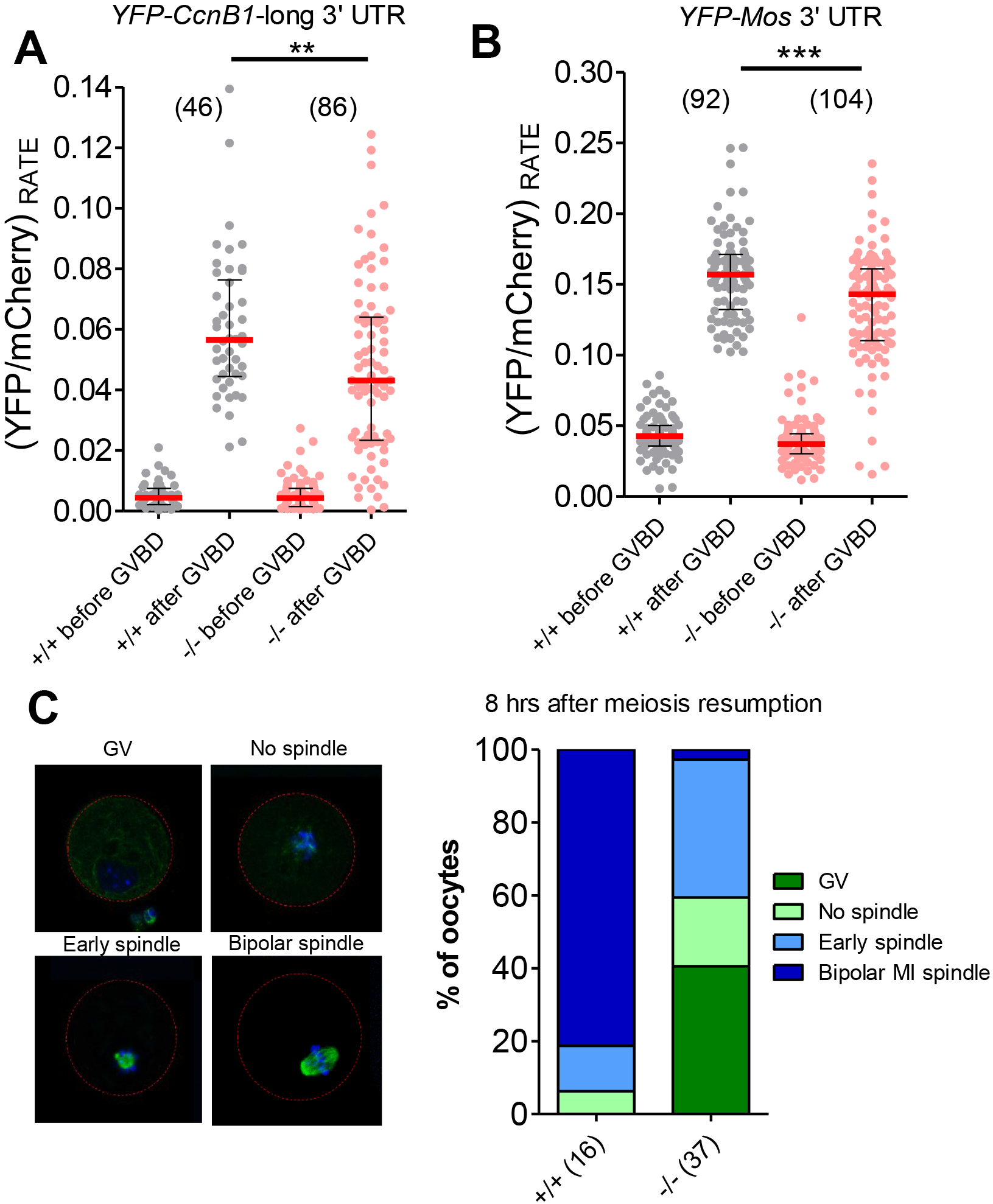
MI spindle formation and activation of *CcnB1* and *Mos* translation are disrupted in *CcnB2^−/−^* out mice. **A-B)** Oocytes were injected with a 1:1 mix of mCherry-polyadenylated and either *YFP-CcnB1-long* 3’UTR **(A)** or *YFP-Mos* 3’UTR **(B)**. After overnight incubation, oocytes were release in Cilostamide-free medium, and brightfield, YFP, and mCherry images were acquired every 15 mins for 25 hrs. YFP signals were normalized by plateaued mCherry signals (YFP/mCherry). The normalized rate of YFP accumulation was calculated before (0-2 hrs) and after (4-6 hrs) GVBD for each singles oocyte. Rates were plotted as the median (red) and interquartile range. T-tests were used to evaluate statistical significance; “**”: p= 0.0058, “***”: p< 0.0001. **C)** Oocytes were released in Cilostamide-free medium and fixed 8 hrs after meiotic resumption. The spindle and the chromatin were visualized with β-tubulin 488 antibody, and DAPI, respectively. Representative pictures are shown for oocytes arrested in prophase I, GVBD without a spindle, early spindle, and bipolar MI spindle. Oocytes were scored for maturation stage and plotted as percentage of oocytes at each stage. Number of oocytes in each group is reported.

It is well established that spindle formation requires protein synthesis and, in particular, the accumulation of CCNB1, which is necessary to increase MPF activity (Davydenko et al., 2013; Hampl and Eppig, 1995; Winston, 1997). Since we have shown that *CcnB*2^−/−^ oocytes have less pre-MPF activity (Fig. 3 B-C) and that the rates of YFP-CCNB1-long 3’UTR accumulation are decreased (Fig. 5 A), we predicted a delay in the time of spindle formation in *CcnB*2^−/−^ oocytes. *CcnB*2^+/+^ and *CcnB*2^−/−^ oocytes were maturated *in vitro* and fixed eight hours after meiotic resumption and the spindle was visualized via β-tubulin staining. While more than 80 percent of *CcnB*2^+/+^ oocytes have a MI bipolar spindle, 60 percent of the *CcnB*2^−/−^ oocytes display no or early spindle (Fig. 5 C). Together, these findings support a role of CCNB2 in CDK1-dependent translation of CCNB1 and MOS and the timely assembly of MI spindle.

### Delayed MI/anaphase I transition in CcnB2^−/−^ oocytes is associated with defective APC activity and persistent activation of the SAC

To define whether additional defects in meiotic progression are associated with CCNB2 depletion, we examined the ability of *CcnB*2^−/−^ oocytes to complete meiosis I. First polar body extrusion (PBE I) is both significantly delayed and decreased in these oocytes (Fig 6 A). In addition, only 30 percent of the *CcnB*2^−/−^ oocytes reach MII with a well formed spindle and aligned metaphase chromosomes, while the rest of the *CcnB*2^−/−^ oocytes are arrested in MI or at telophase I (Fig. 6, B and C). This phenotype is not due to unfavorable *in vitro* culture conditions because the same analysis of *in vivo* ovulated oocytes also clearly shows compromised progression to MII in *CcnB*2^−/−^ oocytes (Fig. S5 A).

**Fig. 6.**
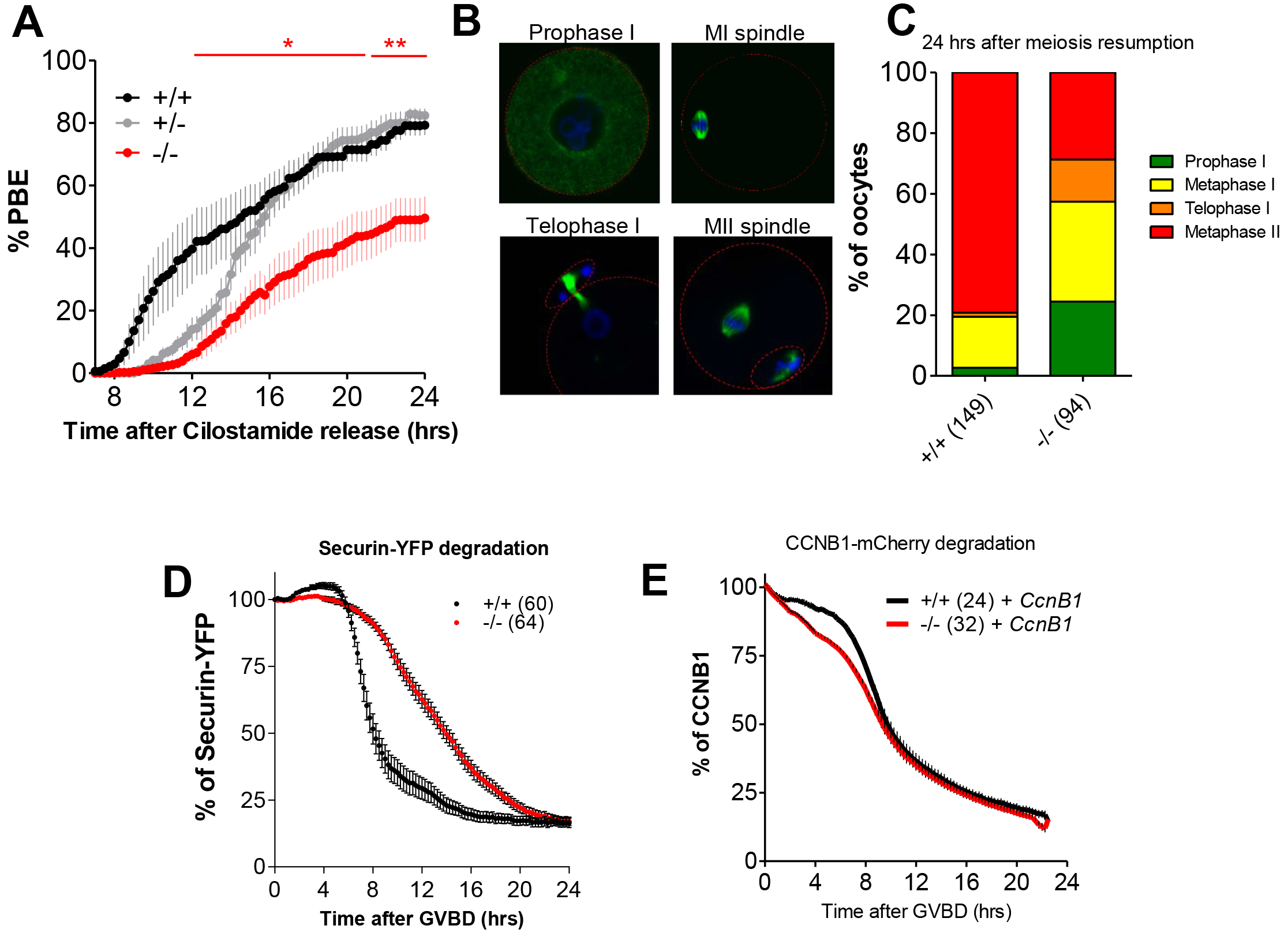
A population of *CcnB*2^−/−^ oocytes fails to complete meiosis I because of altered APC activation. Oocytes were released in Cilostamide-free medium and brightfield images were captured every 15 mins. **A)** The cumulative times of PBE were plotted and t-tests between *Ccnb*2^+/+^ and *CcnB*2^−/−^ oocytes performed (red asterisks); “*”: p< 0.05, “**”: p< 0.01. **B-C)** Oocytes were released in Cilostamide-free medium and were fixed after 24 hrs. The spindle and the chromatin were visualized with β-tubulin 488 antibody, and DAPI, respectively. **B)** Representative pictures are shown for oocytes arrested in prophase I, MI, telophase I, and MII. **C)** Oocytes were scored for maturation stage (reported in panel B) and plotted as percentage of oocytes at each stage. **D)** Oocytes were injected with mRNA encoding the APC substrate *Securin-YFP* and, after 17 hrs incubation, released in Cilostamide-free medium. Securin-YFP level was measured every 15 mins. **E)** Oocytes were injected with mRNA encoding for *CcnB1-mCherry* and, after 3 hrs incubation, released in Cilostamide-free medium. CCNB1-YFP levels were measured every 15 mins.

A compromised MI/anaphase I transition may result from defects in the activation of the APC/CDC20 complex, which promotes the degradation of securin and cyclins. To assess this possibility, oocytes were injected with *Securin-YFP* mRNA to monitor the kinetics of APC activation in live oocytes. Securin degradation is temporally delayed and inefficient in *CcnB*2^−/−^ oocytes (Fig. 6 D). The rate of securin degradation was calculated between six and 10 hours after GVBD and *CcnB*2^−/−^ oocytes display a 50 percent decrease in degradation rates (Fig. S5 B).

It has been reported that CDK1 activates APC directly and indirectly (Adhikari et al., 2014; Golan et al., 2002; Lahav-Baratz et al., 1995; Qiao et al., 2016; Yang and Ferrell, 2013), and this activity is critical for satisfaction of the spindle assembly checkpoint (SAC) (Lara-Gonzalez et al., 2012). Using a FRET probe, we measured changes in CDK1 activity of single oocytes between two and six hours after GVBD. *CcnB*2^−/−^ oocytes that extrude the first polar body have increased CDK1 activity similar to that of *CcnB*2^+/+^ oocytes, while CDK1 activation is decreased in *CcnB*2^−/−^ oocytes unable to complete meiosis I (Fig. S5 C). Moreover, APC activation is rescued by overexpression of CCNB1 in *CcnB*2^−/−^ oocytes (Fig. 6 E, Fig. S5 D).

Since a population of *CcnB*2^−/−^ oocytes is unable to complete meiosis I, we investigated if the inability to progress to anaphase I may be due to the presence of unattached chromosomes and an active SAC. MAD2 co-localization with the kinetochore, as visualized by CREST antibody, has been used as an tool to measure SAC activation (Collins et al., 2015; Gui and Homer, 2012). *CcnB*2^+/+^ and *CcnB*2^−/−^ oocytes were fixed at either seven or 24 hours post-meiotic resumption and MAD2/CREST ratios were measured. *CcnB*2^+/+^ oocytes display low MAD2 signals on the kinetochores at both times, indicating that the SAC had been satisfied (Fig. 7, A and B). Pharmacological depolymerization of the spindle with Nocodazole strongly activates the SAC in *CcnB*2^+/+^ oocytes, and MAD2 localizes on the spindle at most of the kinetochores (Fig. 7, A and B). *CcnB*2^−/−^ oocytes that reach MII display MAD2 levels on the kinetochores that are comparable to that of *CcnB*2^+/+^ oocytes arrested in MII (Fig. 7, A and B). Conversely, *CcnB*2^−/−^ oocytes unable to complete meiosis I after 24 hours display significantly higher levels of MAD2 on the kinetochores than *CcnB*2^++^ MI oocytes (Fig. 7, A and B). This finding suggests that the MI-arrested *CcnB*2^−/−^ oocytes do not transition to anaphase I because SAC is still active. It is known that oocytes can tolerate some unattached kinetochores and still proceed to anaphase I (Lane et al., 2012); therefore an additional experiment was performed to confirm that an active SAC is indeed the cause of the MI arrest in *CcnB*2^−/−^oocytes. Inhibition of MPS1 with Reversine is known to prevent MAD2 localization on the kinetochores and suppresses the activity of SAC (Tipton et al., 2013). The length of meiosis I was measured as the time interval between GVBD and PBE I (Fig. 7 C). Meiosis lasts longer in *CcnB*2^−/−^ oocytes than in *CcnB*2^+/+^ oocytes (WT: 8.25 ±2.5 hrs; KO: 11.7 ±3.0 hrs). Pharmacological SAC inhibition shortens meiosis I in both *CcnB*2^+/+^ (5.6 ±0.8 hrs) and *CcnB*2^−/−^ (5.3 ±1.1 hrs) oocytes, and virtually all the oocytes complete meiosis I with comparable time courses, regardless of the genotype. Thus, the persistent SAC activity in *CcnB*2^−/−^ oocytes is indeed the cause of the delayed in PBE I timing and/or the arrest in MI. All together, these findings indicate that *CcnB*2^−/−^ oocytes are less efficient in satisfying the SAC, leading to defective APC activation and, ultimately, a delayed or failed to exit from MI. These defects are rescued by overexpression of CCNB1.

**Fig. 7.**
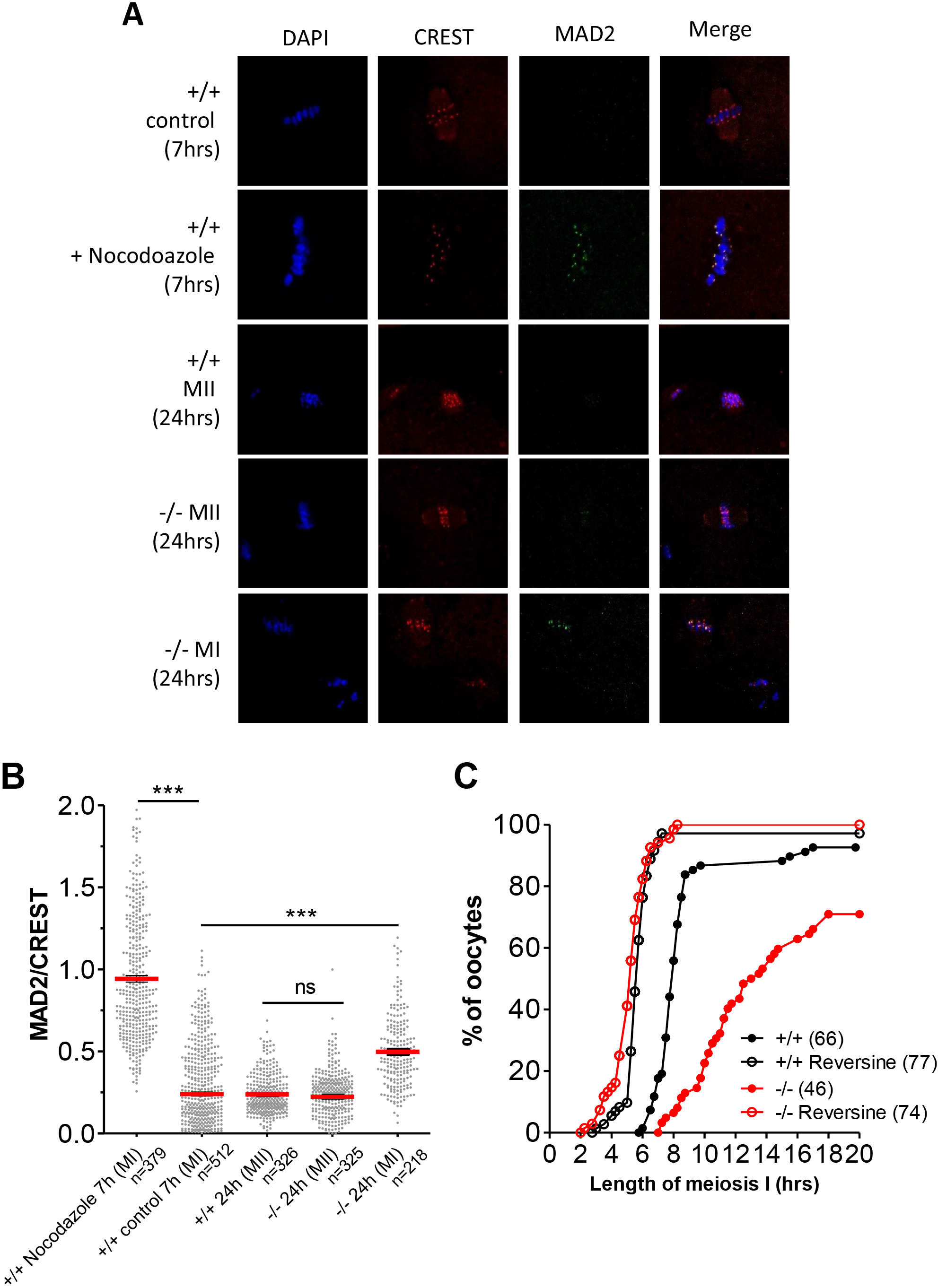
A population of *CcnB*2^−/−^ oocytes arrests in MI because of persistent SAC activity. **A-B)** Oocytes were released in Cilostamide-free medium and fixed at indicated times. Where specified, oocytes were treated with Nocodazole 15 mins before fixation. MAD2, CREST, and chromatin were visualized with specific antibodies and DAPI, respectively. **A)** Representative pictures are shown for each condition. **B)** The amount of MAD2 localized at each single kinetochore was quantified by measuring the ratio between MAD2 and CREST. Number of kinetochore analyzed is reported below each scatter plot. T-tests were used to evaluate statistical significance; ns: “not significant,” “***”: p< 0.0001. **G)** Oocytes were released in the absence or presence of 100 nM Reversine. Time of GVBD and PBE was determined through brightfield images acquired every 15 mins for 24 hrs. The length of meiosis I was calculated as the time between GVBD and PBE.

## DISCUSSION

Our findings conclusively establish that, in the oocyte, CCNB2 plays a critical role during meiosis I both at the G_2_/M and the MI/anaphase I transitions — functions that are not compensated for by endogenous CCNB1. CCNB2 contributes to pre-MPF activity during prophase and is required to generate sufficient MPF to progress through maturation in a timely and efficient fashion (Fig. 3). Moreover, loss of CCNB2 is associated with ovulation of immature oocytes and/or oocytes with compromised developmental competence, resulting in decreased fecundity in *CcnB*2^−/−^ females (Fig. 2).

Initial evidence suggests that *CcnB2* mRNA is translated at a higher rate than *CcnB1* mRNA in prophase. First, the overall levels of *Ccnb2* and *Ccnb1* transcripts are comparable, *CcnB2* is translated at a higher rate than *CcnB1* in GV-arrested oocytes (Fig. 1, A and B). Second, two *CcnB2* and three *CcnB1* isoforms with 3’ UTRs of varying lengths are expressed. While both *CcnB2* isoforms are highly translated in prophase, only the short isoform of *CcnB1* is translated at high levels during this time (Fig. 1 D) (Han et al., 2017; Yang et al., 2017). Third, our previous polysomal array data confirms higher recruitment of *CcnB2* to the polysome as compared to *CcnB1* (Han et al., 2017). Furthermore, we have previously reported similar rates of degradation for the two proteins in GV oocytes in the presence of cycloheximide (Han et al., 2017). Taken together, these data indicate that CCNB2 protein is present at higher concentrations than CCNB1 in GV-arrested oocytes, supporting a central role for CCNB2 during meiosis I.

CCNB2 is critical to generate sufficient levels of pre-MPF in the GV oocyte, and by using both whole cell and *in vitro* kinase assays, we show that *CcnB*2^−/−^ oocytes have decreased CDK1 activity as compared to *CcnB2^+/+^* oocytes (Fig. 3 B, Fig. S2 B). Due to this decreased pre-MPF activity, conversion of pre-MPF to MPF is also affected. This was confirmed by measuring MPF activation via a FRET probe (Fig. S3, G and H) and by observing the export kinetics of WEE1B from the nucleus, an event shown to be CDK1 dependent (Fig. 4 B) (Oh et al., 2010). CDK1 activation in *CcnB*2^−/−^ oocytes no longer displays a switch-like property as in *CcnB*2^+/+^, but rather increases slowly over a long period of time. As a consequence of this gradual increase, GVBD becomes an inefficient process, becoming error-prone (Fig. 3 D). Thus, a subset of *CcnB*2^−/−^ oocytes are unable to transition from prophase I to MI even after prolonged culture times.

Previous attempts to deplete oocytes of CCNB2 with MO indicate an 80 percent decrease in meiotic reentry three hours post-IBMX release (Gui and Homer, 2013). This finding led the authors to conclude that CCNB2-depleted oocytes do not reenter meiosis. Our data, instead, show that meiotic reentry is delayed, but not abolished. A possible explanation of these distinct outcomes is the distinct effects of acute and chronic depletion of CCNB2 (El-Brolosy and Stainier, 2017). It should be pointed out that in the study of Gui and Homer, only the first three hours of meiotic resumption were reported, and therefore, any further delay in maturation may have been overlooked. Similarly, Li et al. observed a decrease in meiotic maturation in *CcnB*2^−/−^ oocytes. Again, only the first three hours of maturation were reported. Therefore, neither studies explored CCNB2 function beyond the G_2_/M transition.

In our study, we have further surveyed the role of CCNB2 throughout meiotic maturation and have detected additional phenotypes. Indeed, depletion of CCNB2 disrupts the increase in CDK1 activity that normally occurs during prometaphase (Fig. S5 C). This defective CDK1 activation may be dependent on both direct effects due to the absence of CDK1/CCNB2 complex and indirect effects on the activity of CDK1 in complex with CCNB1. This is predicted by the decreased rate of *CcnB1* mRNA translation in a subset of *CcnB*2^−/−^ oocytes (Fig. 5). Additionally, the decreased translation of *Mos* mRNA likely affects the positive feedback between the ERK pathway and MPF (Nebreda and Ferby, 2000)(Fig. 5).

Deficient CDK1 activity in MI has several consequences. There is a delay in spindle assembly (Fig. 5C), which recapitulates previous experiments using pharmacological inhibition of CDK1 (Davydenko et al., 2013). Similarly, SAC inactivation is incomplete in the *CcnB*2^−/−^ oocytes leading to defective APC activation. Indeed, stable microtubule attachments to the kinetochores have been shown to depend on the increase in CDK1 activity (Davydenko et al., 2013). In agreement with these findings, disruption of CDK1 activity via depletion of CCNB2 results in persistent MAD2 loading onto the kinetochores (Fig. 7 B). Inhibition of the checkpoint with Reversine restores the oocyte ability to progress to anaphase (Fig. 7 C). Downstream of SAC satisfaction is the activation of APC/C. Our data indicate that, without CCNB2, there is a major delay in APC activation as measured by securin degradation (Fig. 6 D). In *CcnB*2^−/−^ oocytes, APC activation is not switch-like as in *CcnB*2^+/+^ oocytes, but instead, prolonged and gradual—in agreement with the idea that an threshold of CDK1 activity is required for full APC activation (Yang and Ferrell, 2013). Thus, in *CcnB*2^−/−^ oocytes, the switch-like entry into and exit from meiosis I is disrupted, resulting in non-synchronous, delayed, and sometimes failed G_2_/M and MI/anaphase I transitions.

In summary, our findings establish a unique function for CCNB2 during mouse oocyte meiosis that cannot be compensated for by endogenous levels of CCNB1. We show, however, that overexpression of exogenous CCNB1 completely rescues the effect of CCNB2 depletion both during the GV/MI (Fig. 3 E) and the MI/anaphase I transitions (Fig. 6 E). In a reciprocal study using *CcnB1^−/−^* mice, endogenous CCNB2 alone is able to drive oocyte progression through meiosis I, but the oocytes could not progress to MII. However, overexpression of exogenous CCNB2 rescues the MII entry (Li et al., 2018, 2). Taken together, these two complementary studies indicate that the two cyclin proteins have overlapping function at the molecular level. The difference in ability of exogenous and endogenous proteins to rescue meiosis progression is likely due to the constitutive overexpression of the exogenous protein, while expression of the endogenous protein is under the temporal control of the oocyte translational program. Indeed, we have previously shown that the translation of *CcnB2* and *CcnB1* mRNAs is markedly different; *CcnB2* is constitutively translated in GV and MI, but decreased in MII, whereas the recruitment of *CcnB2* mRNA to the polysome increases at MI and further increases at MII (Fig. 1) (Han et al., 2017). Translation of endogenous *CcnB2* decreases in MII explaining its inability to compensate for the absence of CCNB1 at this stage. These findings emphasize the importance of the post-transcriptional program in regulating meiotic divisions. Indeed, the distinct temporal translational control of these two related genes dictate the appropriate cyclin levels to orchestrate faithful progression through meiosis.

Given our findings, we propose that the CCNB2-depleted oocytes may be used as a model for defective CDK1 activation throughout meiosis, a condition that may be a significant cause of meiotic maturation block and infertility in humans.

## ACKNOWLEDGEMENTS

We acknowledge Dr. Tim Hunt and Dr. Jonathon Pines for sharing the *CcnB2^−/−^* mice and Dr. Rey-Huei Chen for the gift of the MAD2 antibody. The authors are indebted to Dr. Sophie Dumont for the helpful discussion and advice on the SAC measurements. These studies were supported by NIH R01 GM097165 and GM116926 to MC. EMD is supported by a fellowship from the Lalor Foundation.

## Materials and methods

### Mice, oocyte collection, and microinjection

All experimental procedures involving mouse were approved by the Institutional Animal Care and Use Committee of UCSF (Protocol: AN101432). C57BL/6 female mice (21-24 days) were primed with 5 units of PMSG and were sacrificed 44-48 hours later to collect GV-arrested oocytes. For collection of MII-arrested oocytes, females were primed with 5 units of PMS, after 48 hrs, injected with hCG, and after 13 hrs, sacrificed for egg retrieval. Cumulus-enclosed oocytes from antral follicles were isolated, and mechanically denuded in HEPES modified Minimum Essential Medium Eagle (Sigma-Aldrich, M2645) supplemented with 1μM Cilostamide (Calbiochem, 231085). When specified, oocytes were microinjected with 5-10 pl of mRNA. Oocytes were then cultured at 37°C with 5% CO_2_ in MEM-α medium (Gibco, 12561-056) supplemented with 0.2 mM sodium pyruvate, 75 μg/ml penicillin, 10 μg/ml streptomycin, 3 mg/ml bovine serum albumin (BSA), and 1 μM Cilostamide for 3 hrs or 16 hrs as indicated in the figure legends.

### Plasmid construct and mRNA preparation

(C483S)-CDC25B and (K237A)-WEE1B coding sequence were cloned upstream of the YPet coding sequence. The CCNB1 and CCNB2 open reading frame sequences were obtained by sequencing oocyte cDNA and cloned upstream of the mCherry coding sequence. The *CcnB1, CcnB2*, and *Mos* 3’UTR sequences were also obtained in the same manner and cloned downstream of the YPet coding sequence. All constructs were prepared in the pCDNA 3.1 vector containing a T7 promoter and fidelity was confirmed by DNA sequencing. mRNA of all the reporters were *in vitro* transcribed with mMESSAGE mMACHINE T7 Transcription Kit (Ambion, AM1344); when specified, polyadenylation was achieved using Poly(A) Tailing Kit (Ambion, AM1350). All the messages were purified using MEGAclear Kit (Ambion, AM1908). mRNA concentrations were measured by NanoDrop and message integrity was evaluated by electrophoresis.

### Time-lapse microscopy, analysis of protein translocation, and *YFP*-3’UTR translation

Time-lapse experiments were performed using a Nikon Eclipse T2000-E equipped with mobile stage and environmental chamber to 37°C and 5% CO_2_. Filter set: dichroic mirror YFP/CFP/mCherry 69008BS; for Ypet channel (Ex: S500/20x 49057 Em: D535/30m 47281), mCherry channel (Ex: 580/25x 49829 Em: 632/60m). (C483S)-CDC25B-YFP, (K237A)-WEE1B-YFP, securin-YFP, or CCNB1-mCherry were injected at 300 ng/μL. After injection, oocytes were incubated for 16 hrs to allow expression of the probes. Ratios of nuclear and total probe were calculated after subtraction of the background. Rate of translocation were calculated as the slope of the line obtained by linear regression. *YFP*-3’UTR reporters we co-injected with polyadenylated *mCherry* at 12.5 μg/μL each. After injection, oocytes were incubated for 16 hrs to allow expression of the probes. YFP signals were normalized by the plateaued level of mCherry signal to control of amount of injection. Rates were calculated with YFP/mCherry ratios as the slope of the curve obtained by linear regression of the time points indicated.

### RiboTag-Immunoprecipitation and RNASeq

Oocytes were collected in minimal volumes (5-10 μl) in 0.1% polyvinylpyrrolidone (PVP) in DPBS, flash frozen in liquid nitrogen, and stored at −80°C. Samples were thawed, randomly pooled to yield a total of 200 oocytes per time point per replicate, and 300 μl supplemented homogenization buffer (sHB) was added to each pooled sample. The homogenates were then vortexed for 30 secs, flash frozen in liquid nitrogen, and allowed to thaw at room temperature (RT); this was repeated twice. Finally, the homogenates were centrifuged for 10 mins at maximum speed at 4°C and the supernatant (IP soup) was collected in new tubes. Meanwhile, the appropriate volume (50 ul per sample) of Dynabeads™ Protein G (Invitrogen) was washed three times in 500 μl homogenization buffer (HB) on a rotor at 4°C for 5 mins per wash. An additional two washes were performed with 500 ul sHB on a rotor at 4°C for 10 mins per wash. The final wash solution was removed and the beads were eluted in the original volume of sHB and kept on ice. 20 μl cleaned beads was added to each IP soup to pre-clear on a rotor at 4°C for 1 hr. The beads were removed via a magnetic rack and 15 μl of IP soup was collected from each sample in 200 ul of RLT buffer (Qiagen) to serve as the input samples. Input samples were frozen and kept at −80°C until RNA extraction. 3 ul (3 ug) anti-HA.11 epitope tag antibody (901501, BioLegend) was added to each of the remaining IP soups and all samples were incubated on a rotor at 4°C for 4 hrs. 30 ul clean beads were then added to the samples and incubated overnight on a rotor at 4°C. The beads (now bound by HA-tagged ribosomes and the associated mRNAs) were washed five times in 1 ml of wash buffer with 1 M urea (uWB) on a rotor at 4°C for 10 mins per wash. The final uWB wash was removed and 250 μl RLT buffer was added to each sample and vortexed for 30 secs. RNA extraction was performed following the Rneasy Plus Micro Kit protocol (Qiagen). Samples were eluted in 10 μl of RNAse-free water. RNA samples were sent to the Gladstone Institutes Genomics Core for quality control using Bioanalyzer (Agilent) and cDNA library preparation with the Ovation RNA-Seq System V2 (NuGen). Samples were sequenced using the HiSeq400 platform.

### Western blot

Oocytes were collected in 0.1% PVP in DPBS and then boiled for 5 min at 95°C in 1× Laemmli Sample Buffer (Bio-Rad) supplemented with with β-mercaptoethanol. Lysates were resolved in 10% Laemmli gels and transferred onto Supported Nitrocellulose Membranes. Membranes were incubated in the primary antibody overnight at 4°C; Antibodies and dilutions used: CCNB2, 1:1,000 (R&D Systems, AF6004); CCNB1, 1:500 (Abcam, ab72); (β-actin, 1:1,000 (Abcam, ab8227); CDK1, 1:1,000 (Santa Cruz); CPEB1, 1:1,000 (Abcam, ab73287); T320-pp1, 1:30,000 (Abcam, ab62334); GST, 1:10,000 (Sigma).

### Immunofluorescence

Oocytes were fixed in DPBS supplemented with 0.1% Triton X-100 and 2% formaldehyde (Sigma, 28908) for 30 mins. After three 10 min washes with blocking buffer, the oocytes were incubated overnight in blocking buffer (1x PBS, 0.3% BSA, 0.01% Tween), then permeabilized for 15 mins in DPBS supplemented with 0.3% BSA and 0.1 % Triton X-100. After three 10 min washes with blocking buffer, oocytes were incubated for one hr in primary antibody diluted in blocking buffer. The antibodies used: β-tubulin-488, 1:100 (Cell Signaling Technology, 3623); CREST, 1:200 (ImmunoVision); MAD2, 1:200 (Dr. Rey-Huei Chen, Academia Sinica, Taipei). After three 10 min washes with blocking biffer, the membrane was incubated for one hr with the appropriate secondary antibody, 1:500 (Alexa-568 goat anti-human; Alexa-488 goat anti-rabbit). Oocytes were washed again for 10 mins, three times in blocking bugger and mounted with VECTASHIELD Mounting Medium with DAPI (Vector, H-1200). Pictures were acquired with a confocal Nikon C1 SI equipped with X60 oil immersion lens.

### *In vitro* CDK1 kinase assay

Oocytes were collected in 30 μl of 2X kinase buffer (100 mM Hepes, 30 mM MgCl_2_, 2 mM EGTA, 10 mM CaCl_2_, 2 mM DTT, 2 μg/ml Leupeptin, 2 μg/ml Aprotinin, 2 μM Okadaic Acid). Oocytes were lysed by freezing and thawing in liquid nitrogen two times. Extracts were incubated at 30°C for 15 mins in presence of 0.1 mM ATP, 10 mM DTT, 2 μM Okadaic acid, and 2 μg of recombinant peptide PP1-GST as the substrate. PP1-GST was produced as previously described (Daldello et al., 2015). Reactions were stopped by adding Laemmli Sample Buffer and boiling at 95°C for 5 mins. CDK1 activity was measured by quantifying the Western blot signal of phosphorylated T320 of the PP1-GST substrate.

### Data processing, quantification, and statistical analysis

Visual quality checks of RNASeq reads were performed using FastQC and reads were them trimmed with Trimmomatic. Alignment of the reads to the mouse genome was performed by Hisat2, .bam files were sorted and indexed using Samtools, and count files were generated by HTSeq. TMM normalization and the remaining RNASeq statistical analyses were done through edgeR. MAD2/CREST signals were quantified with Fiji. T-tests and non-parametric Mann-Whitney tests were performed using the GraphPad Prism 7.

## Supplementary methods

### FRET experiment

The CDK1 FRET sensor (2327) was a gift from Dr. Jonathon Pines (Addgene, 26064). Oocytes were injected with 5-10 pl of FRET sensor mRNA at 300 ng/μl, and, after 16hrs incubation, fluorescence level was quantified as described in the methods section. Intensity signals of YFP/YFP, CFP/CFP and CFP/YFP channels were subtracted by the background. The CFP/YFP channel was corrected for the YFP bleed-through and FRET was calculated as (CFP/YFP)/(CFP/CFP).

**Fig. S1.**
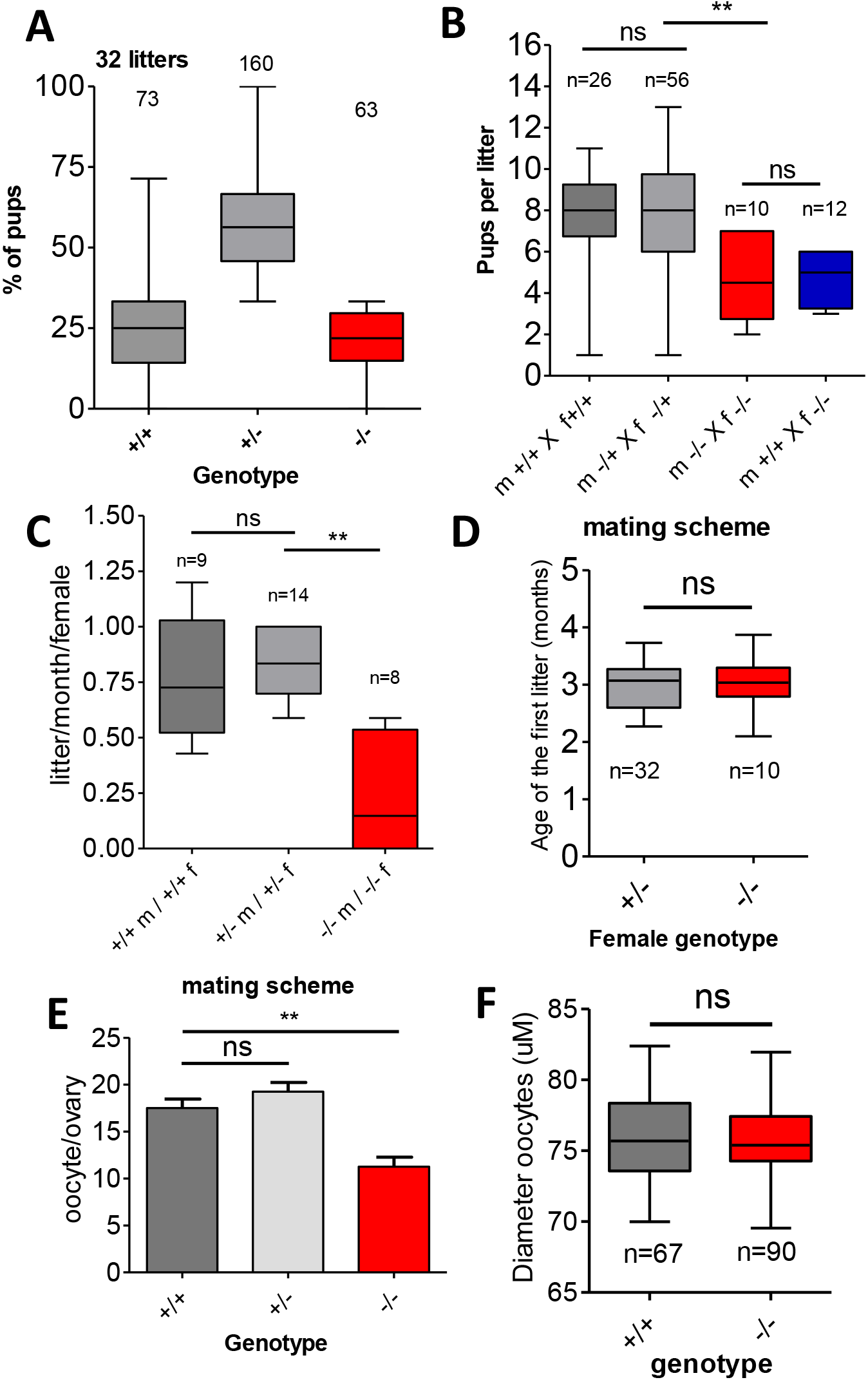
**A)** Pups from +/−♂ × +/−♀ matings were genotyped and classified according to their genotype. Thirty-two litters from 12 mating couple were analyzed. **B)** Number of pups per litter was recorded. The number of litters analyzed is displayed. T-tests were performed to evaluate the statistical significance; “ns”: not significant, “**”: p= 0.0014. **C)** The frequency of parturition was measured for different mating schemes and expressed as number of litters per month per female. T-tests were performed; “ns”: not significant, “**”: p= 0.0022.**D)** Age at the first litter for different mating schemes. Breeding was initiated when animals reached four weeks of age. **E)** Number of oocytes retrieved from the ampullae per ovary after PMSG and hCG treatment. **F)** Diameter of the oocytes was measured by inspecting brightfield recordings.

**Fig. S2.**
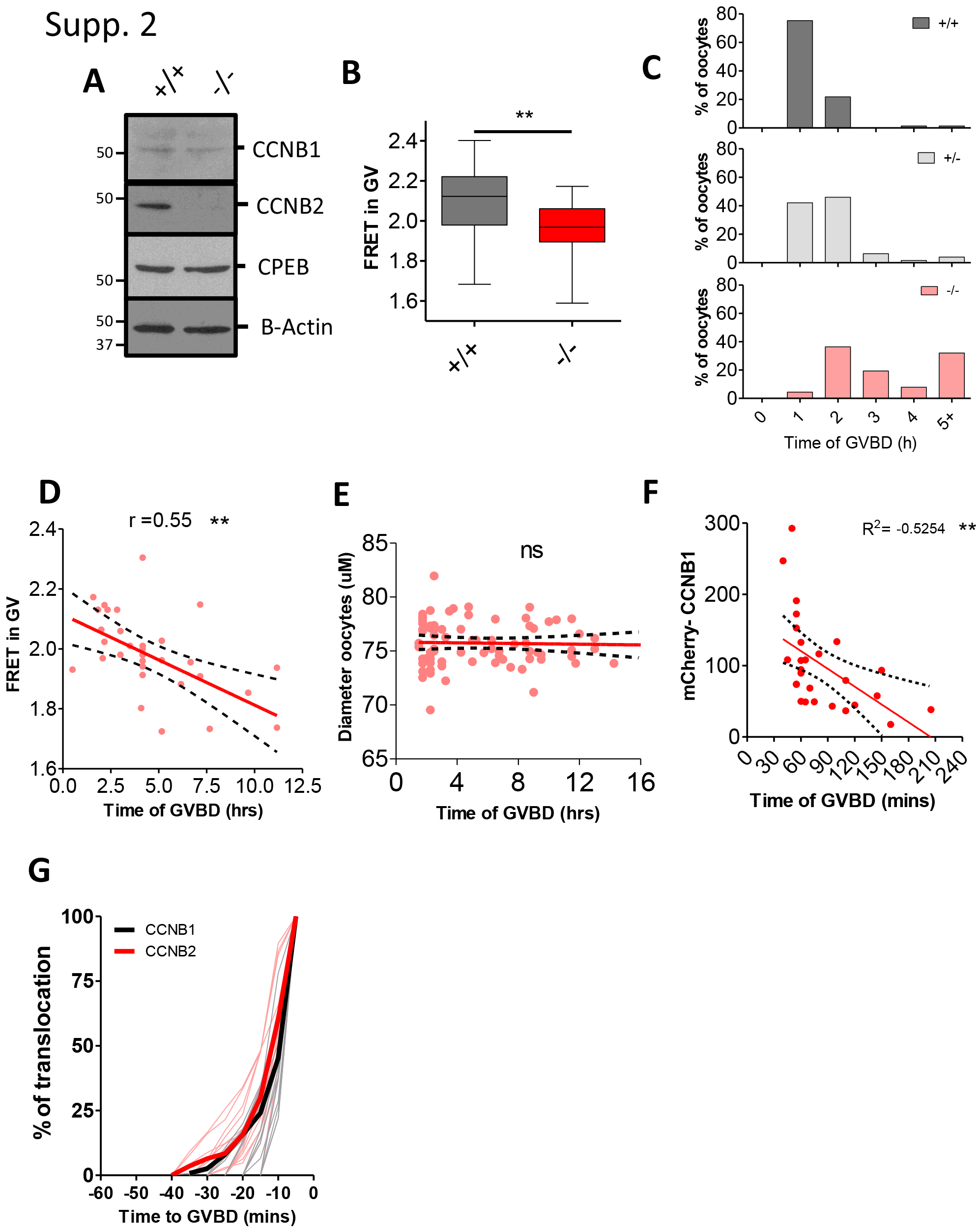
A) Western blot analysis of 150 oocytes from *CcnB*2^+/+^ and ^−/−^ mice. **B)** Oocytes were injected with a probe specific for CDK1. After 18 hrs, four frames were recorded in brightfield, YFP/YFP, CFP/CFP, and CFP/YFP. FRET is expressed as (CFP/YFP)/(CFP/CFP); “**”: p= 0.001. **C)** Time of GVBD from Fig. 3 B was replotted as a histogram to highlight the presence of two different populations in the *CcnB*2^−/−^ oocytes. **D)** The level of FRET in GV-arrested oocytes was correlated the GVBD time of each oocyte. The Pearson coefficient (p= −0.55) has been calculated with its associated p-value; “**”: p= 0.0013. **E)** The diameter of *CcnB*2^−/−^ oocytes was correlated to GVBD time; “ns”: not significant. **F)** Level of expression of mCherry-CcnB1 in each oocyte was correlated with GVBD time. Pearson coefficient (p=−0.53) has been calculated with its associated p value; “**”: p= 0.0049. **D-F)** The best-fit line is displayed in red and the 95 percent interval of confidence is represented as dashed lines.

**Fig. S3.**
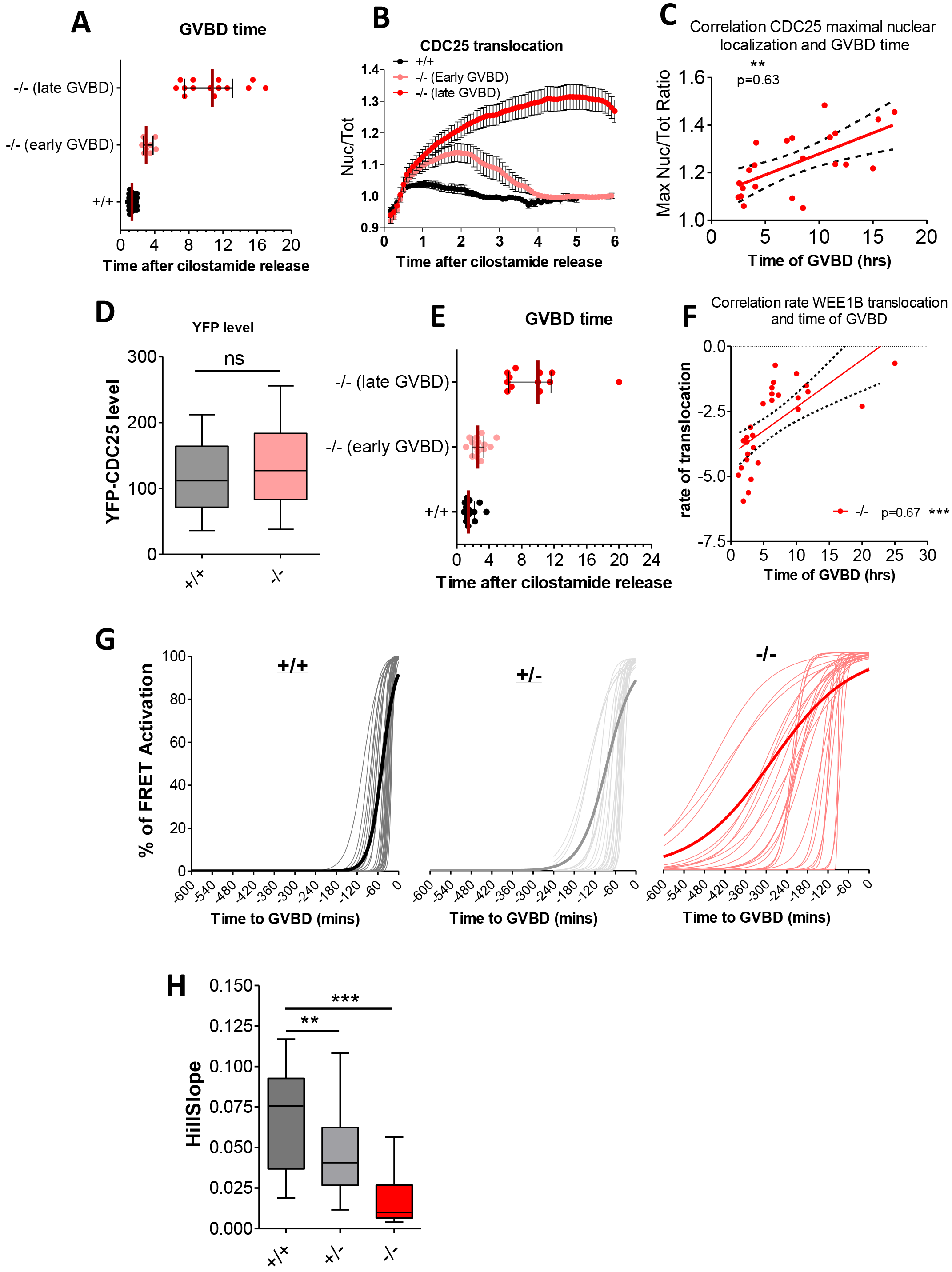
**A)** GVBD time of oocytes from Fig. 4, A and C was determined by the inspection of the brightfield recordings. **B)** The average of the nuclear/cytoplasmic ratios of CDC25B-YFP signals that was used to calculate the rates Fig. 4, A and C. are plotted. **C)** The maximum nuclear/cytoplasmic ratio of CDC25B-YFP localization of each oocyte was correlated to the oocyte GVBD time. The best-fit line is displayed in red and the 95 percent interval of confidence is represented as dashed lines. The Pearson coefficient (p= 0.63) was calculated with its associated p-value; “**”: p= 0.0016. **D)** The level of the CDC25B-YFP at the start of recording from Fig. 4, A and C was quantified. **E)** GVBD time of oocytes from Fig. 4, B and D was determined by inspection of brightfield recordings. **F)** The rate of WEE1B-YFP translocation of oocytes from Fig. 4, B and D was correlated to GVBD time of the oocytes. The best-fit line is displayed in red and the 95 percent interval of confidence is represented as dashed lines. The Pearson coefficient (p= 0.63) has been calculated with its associated p-value; “**”: p= 0.0016. **G)** *Ccnb*2^+/+^, ^+/−^, and ^−/−^ oocytes were injected with a CDK1-FRET reporter and, after 16 hrs incubation, released in Cilostamide-free medium. YFP/YFP, CFP/CFP and CFP/YFP signals were recorded every 5 mins. Individual FRET time courses were fitted to a sigmoid equation 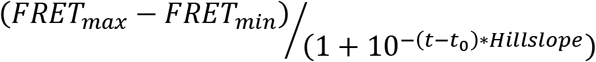 Standardized on the individual GVBD times. Single oocyte time courses are shown as thin lines, while the average of the fitted curves are shown as bolded lines. **H)** Hillslopes from G) were plotted with the median and interquartile ranges. T-tests were used to evaluate statistical significance; “**”: p= 0.0038, “***”: p< 0.0001.

**Fig. S4.**
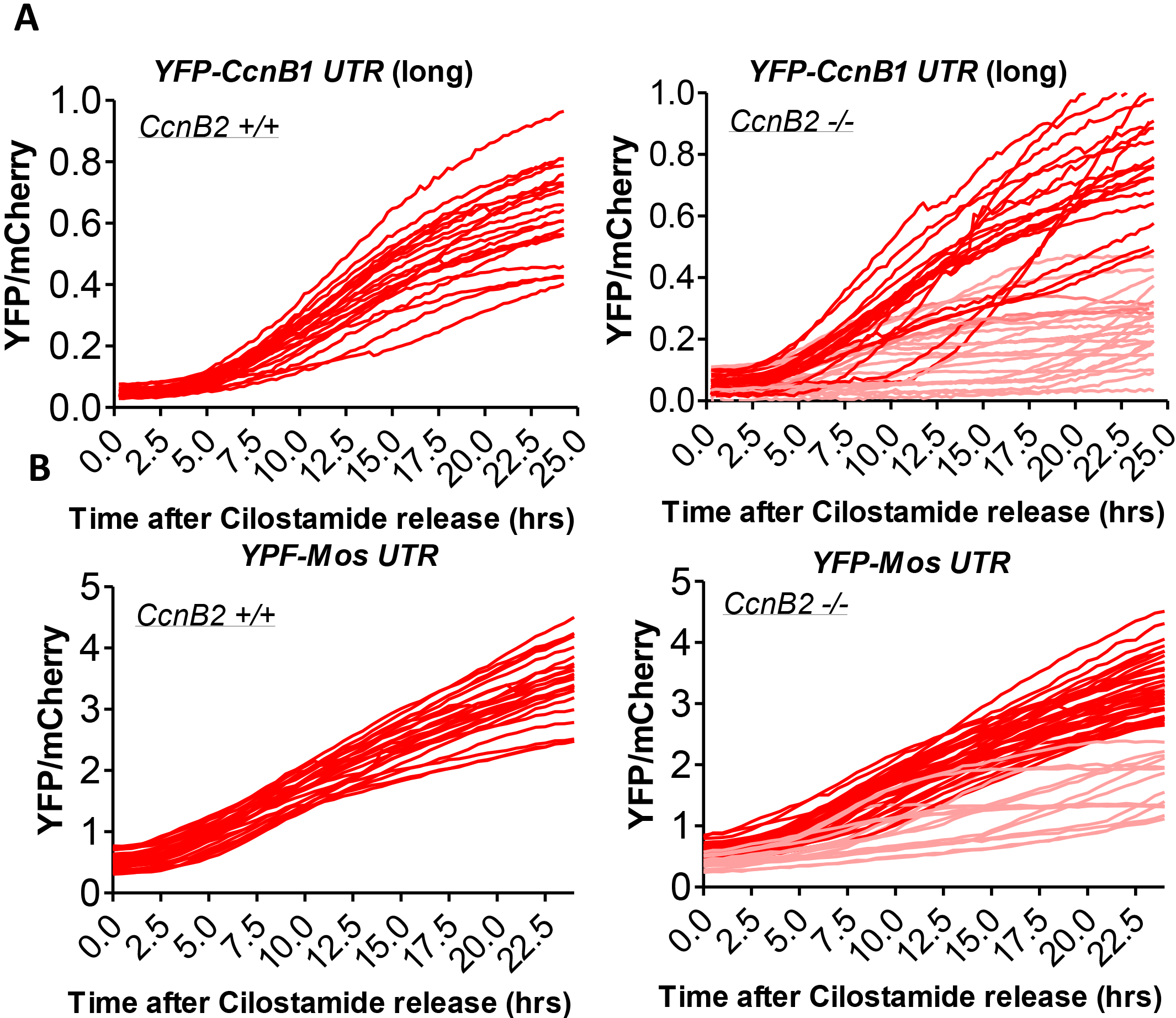
Complete single time courses of *YFP-CcnB1* 3’UTR (Long) **(A)** and *YFP-Mos* 3’UTR **(B)** used to calculate rates in Fig. 5, A and B, respectively.

**Fig. S5.**
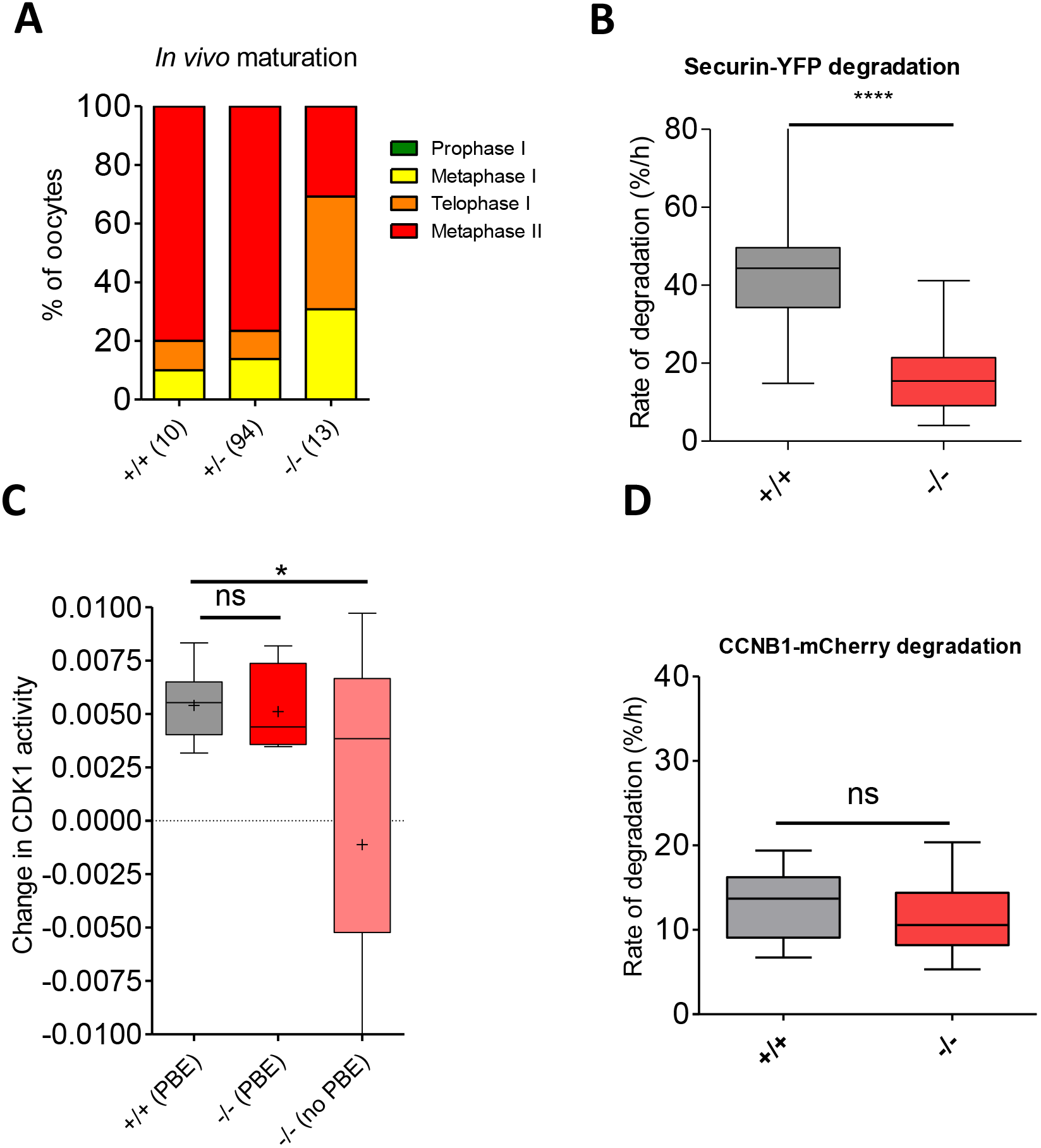
**A)** Oocytes were retrieved after 13 hrs after hCG injection from the ampulla. Oocytes were scored based on the categories illustrated in Fig. 6 B. **B)** The rate of securin degradation was calculated for each oocyte and plotted as the median with the 25/75 intervals of confidence. T-test were used to evaluate statistical significance; “****”: p= 0.0001. C) *Ccnb*2^+/+^ and *^−/−^* oocytes were injected with a CDK1-FRET reporter and, after 16 hrs incubation, released in Cilostamide-free medium. YFP/YFP, CFP/CFP, and CFP/YFP signals were recorded every 15 mins. The rate of change of FRET activity was calculated as the slope of FRET change with a window between 3-6 hrs after GVBD; “*”: p= 0.0011. **D)** The rate of CCNB1 degradation was calculated for each oocyte and plotted as the median with the 25/75 intervals of confidence. T-test was used to evaluate statistical significance; “ns”: not significant.

